# RAVER1 interconnects lethal EMT and miR/RISC activity by the control of alternative splicing

**DOI:** 10.1101/2023.06.14.544976

**Authors:** Alice Wedler, Nadine Bley, Markus Glaß, Simon Müller, Laura Schian, Kingsley-Benjamin Obika, Marcell Lederer, Claudia Misiak, Tommy Fuchs, Marcel Köhn, Roland Jacob, Tony Gutschner, Christian Ihling, Andrea Sinz, Stefan Hüttelmaier

## Abstract

The RAVER1 protein was proposed to serve as a co-factor in guiding the PTBP-dependent control of alternative splicing (AS). Whether RAVER1 solely acts in concert with PTBPs and how it affects cancer cell fate remained elusive. Here we provide the first comprehensive investigation of RAVER1-controlled AS in cancer cell models and reveal a pro-oncogenic role of RAVER1 in tumor growth. This unravels that RAVER1 guides AS in synergy with PTBPs but more prominently serves PTBP1-independent roles in splicing. In cancer cells, one major function of RAVER1 is the control of proliferation and apoptosis, which involves the modulation of AS events within the miR/RISC pathway. Associated with this regulatory role, RAVER1 antagonizes lethal, TGFB-driven epithelial-mesenchymal-transition (EMT) by limiting TGFB signaling. RAVER1-modulated splicing events affect the insertion of protein interaction modules in factors guiding miR/RISC-dependent gene silencing. Most prominently, in all three human TNRC6 proteins, RAVER1 controls AS of GW-enriched motifs, which are essential for AGO2-binding. Disturbance of RAVER1-guided AS events in TNRC6 proteins and other facilitators of miR/RISC activity compromise miR/RISC activity which is essential to restrict TGFB signaling and lethal EMT.

## Introduction

The proper control of splicing, in particular alternative splicing (AS) is central for development and tissue homeostasis by guiding the synthesis of transcript variants ensuring protein isoform diversity (1). In cancer, splicing is largely deregulated due to mutation or aberrant expression of key splicing regulatory proteins (2,3). Previous studies have highlighted the polypyrimidine tract binding protein family (PTBPs) and associated regulatory proteins to serve central roles in the control of AS (4). PTBP1 represses or activates AS via CU-rich motifs (4–6). Association with such motifs via its four RNA recognition motifs (RRMs), oligomerization and the formation of ternary complexes with co-factors was proposed to modulate PTBP-dependent splicing by RNA looping (7,8). Positional rules guiding this PTBP-dependent modulation of splicing were recently demonstrated by a capture RIC-seq (CRIC-seq) method, which enriches RBP-associated RNA-RNA fragments (9).

PTBP-modulated control of AS is frequently disturbed in cancer and deregulated expression of PTBPs is associated with adverse disease outcome in various malignancies (10–12). Elevated expression of PTBP1, the most abundant PTBP paralogue in most cancers, is associated with poor outcome in various solid cancers and associated with clinical properties suggesting roles of PTBPs in cancer progression (12). In agreement, PTBPs have been reported to influence key signaling pathways, e.g. EGFR-, MAPK-, PI3K-signalling, although the molecular mechanisms underlying this regulation remain partially elusive and likely vary in a cancer type-dependent manner (10). A prominent example of PTBP1-dependent control of cancer cell phenotypes is the modulation of a pro-glycolytic shift by PTBP1-dependent enhancement of pyruvate kinase PKM2 and down tuning of PKM1 (13).

Other RNA binding proteins (RBPs) impact on PTBP expression and function (10). The abundance of PTBP1 is controlled in part by nonsense-mediated decay (NMD) regulation, which essentially involves inclusion or skipping of exon 11 in *PTBP1* (14,15). ESPR1 promotes the inclusion of exon 11 leading to increased expression of PTBP1, whereas PTBP1 itself promotes skipping of this exon allowing balanced expression of PTBP1 by negative feed-back control. PTBP-dependent regulation of other splicing events, e.g. AS of tropomyosin (TPM1/2) (16), involves the interaction with other RBPs like RAVER1 (17). This protein was proposed to allow the tethering of distinct PTBPs assembled on the same pre-mRNA at distal and proximal regulatory sites of AS exons to foster out-looping (18). Next to three well-structured N-terminal RNA Recognition Motifs (RRMs), RAVER1 encompasses an extended intrinsically disordered region (IDR), which comprises up to four PTBP-interacting peptide motifs (19,20). These appear essential for the assembly of PTBP-RAVER complexes to direct AS and potentially also promote the association of both proteins in membrane-less protein-RNA condensates in the vicinity of nucleoli, termed perinucleolar compartments (PNC, (17,21)). Intriguingly, PNC size and frequency were proposed to indicate progressed cancer stages and were found associated with adverse disease outcome in cancer (21–23). However, if RAVER1 serves roles in cancer, PNC homeostasis and how this is interconnected with the potentially PTBP-dependent control of splicing in cancer progression remained essentially elusive.

## METHODS

### Plasmids and Cloning

Plasmid generation including vectors, respective templates and restrictions sites were summarized in supplementary table TS1. All constructs were validated by sequencing.

### Cell culture, transfection, and transduction

A549, ES-2, Hep-G2, C-643, PANC-1 cells and HEK293T17 were cultured in DMEM (Gibco), H1650 and H522 cells were cultured in RPMI 1640 (Gibco) and BE(2)-C were cultured in a 1:1 mixture of DMEM/F12 (with HEPES, Gibco) and EMEM (ATCC), all supplemented with 10% FBS and incubated at 37°C with 5% CO_2_. The origin of cancer cell lines is summarized in supplementary Table TS2.

For transfection, 5×10^5^ cells were transfected with 15nM siRNA pools or DNA (Supplementary Table TS1) using Lipofectamine RNAiMax or Lipofectamine 3000 (Thermo Fisher Scientific) according to the manufacturer’s instructions. 24 hours after transfection, cells were transferred in new culture plates. For luciferase studies, 1×10^5^ cells were transfected with 250ng pmirGLO vectors. In overexpression studies, 4×10^6^ cells were transfected with 2µg plasmid using Lipofectamine 3000.

To produce lentiviral vectors, HEK293T17 were transfected with the packaging plasmids psPax2, pMD2.G as well as the lentiviral expression vector pLVX encoding iRFP with Lipofectamine 3000 (Thermo Fisher Scientific) according to manufacturer’s instructions. Lentiviral supernatant was collected 24- and 48-hours post-transfection. Titers were analyzed 72 h post-infection of HEK293T17 cells and determined by flow cytometry (iRFP) using a MACS Quant Analyzer (Miltenyi BioTech). Lentiviral transduction for downstream experiments was accomplished at 10 MOI (multiplicity of infection).

### CRISPR/Cas9 knockout

For CRISPR/Cas9 mediated RAVER1 knockout 5×10^5^ A549 cells were transfected with two sgRNA encoding plasmids (psg_RFP_RAVER1_E×1 and psg_RFP_RAVER1_Ex8) and a Cas9 nuclease encoding plasmid (pcDNA-Cas9-T2A-GFP) using Lipofectamine 3000 according to manufacturer’s protocol. 48-hours post-transfection, cells were sorted for RFP and GFP-positive single cell clones using FACS Melody sorter (BD Bioscience). Knockout was validated by Western blotting.

### Cell proliferation, spheroid growth and anoikis resistance

Analyses of 2D cell proliferation, 3D spheroid growth and anoikis resistance were performed as recently described (24). Approximately 1×10^3^ cells were seeded per 96 well for these studies. Plates were analyzed 5 days post seeding using an Incucyte S3 (Sartorius). Additionally, cell viability was measured using CellTiter Glo (Promega) according to the manufacturer’s protocol. All experiments were performed three times and normalized cell viability was determined by normalization to inputs using the median of at least three independent control analyses, essentially as previously described (25).

### Cell cycle analyses

Cell cycle progression was analyzed 72h post-transfection using propidium iodide staining and flow cytometry (MACSQuant Analyzer, Miltenyi Biotech), as recently described in (25).

### RNA isolation and RT-PCR

Total RNA from cell lines was isolated using TRIzol. RNA concentration was determined by nanodrop (Tecan Sparc), and RNA integrity was monitored on a Bioanalyzer 2100 (Agilent). For cDNA synthesis, 2µg of total RNA served as a template using M-MLV Reverse Transcriptase (Promega) and random hexamer primers following manufacturer’s protocol. Semi-quantitative PCR was performed using OneTaq DNA polymerase (NEB) and gene-specific primers (supplementary table TS3) according to the manufacturer’s protocol. PCR products were separated on 2% agarose gel containing ethidium bromide and visualized under UV light.

### Northern blot

Northern Blot analyses of microRNA expression were performed as previously described (24). Utilized probes are summarized in supplementary table TS4.

### Immunoprecipitation

For immunoprecipitations of SBP-tagged RAVER1 constructs, nuclei were first extracted from cell lysate (1x 10^7^ cells per condition) using SBP-IP buffer (150mM KCl, 50mM TRIS-pH 7.5, 5% (v/v) Glycerol, 5mM EDTA-pH 8, 0.25% (v/v) NP-40) supplemented with 100µg/ml digitonin (Sigma). Cells were lysed for seven minutes at room temperature, following centrifugation for three minutes (4000 rpm). Cell pellets were again resuspended in SBP-IP buffer with digitonin, directly centrifuged and supernatant were discarded. Nuclear extracts were then prepared on ice using SBP-IP buffer. Cleared cell lysates were incubated with Dynabeads MyOne streptavidin beads (Life Technologies) for 2 hours at 4°C. After three washing steps with SBP-IP buffer, protein-protein complexes were eluted by addition of NuPage LDS sample buffer (Thermo Fisher Scientific) with 50mM DTT and incubation at 65°C for 10 min. Protein enrichment were analyzed by Western blotting. Co-immunoprecipitations with following LC-MS/MS Analysis (see below) were essentially performed as described above. Instead of SBP-IP, an SBP-IP-MS-buffer (150mM KCl, 50mM TRIS-pH 7.5, 5% (v/v) Glycerol, 5mM EDTA-pH 8, 0.1% (v/v) DDM) was used. For Co-immunoprecipitations of GFP-AGO2 and HA-constructs, cell extracts were prepared using Co-IP buffer (150mM NaCl, 20 mM TRIS-pH 7.5, 1mM EDTA-pH 8, 0.5% (v/v) Triton). Cleared whole cell lysates were incubated with GFP-antibody and Protein G Dynabeads (Life Technologies) as described above.

### Identification of RAVER1 associated proteins by LC-MS/MS

Pulldown samples (three replicates; see immunoprecipitation above) were prepared for liquid chromatography/tandem mass spectrometry (LC-MS/MS) following a modified FASP (filter-aided sample preparation) protocol (26). Briefly, protein samples were incubated and washed with 8 M urea in 50 mM HEPES (4-(2-hydroxyethyl)-1-piperazineethanesulfonic acid), 10 mM TCEP, pH 8.5. Washing steps were performed using 0.5 mL centrifugal filter units (30-kDa cutoff, Sigma Aldrich, Darmstadt, Germany) and two centrifugation steps (14,000× *g*, 10 min). Afterwards, alkylation of proteins was performed with 50 mM iodoacetamide in 8 M urea, 50 mM HEPES, pH 8.5 at room temperature (20 min in the dark). Samples were washed twice with 50 mM HEPES, pH 8.5 (18,000× *g,* 2 x 10 min) before they were incubated with 1 µg trypsin in 50 mM HEPES, pH 8.5 at 37 °C overnight. After enzymatic digestion, samples were acidified with trifluoro acetic acid (TFA) at 0.5 % (*v/v*) final concentration.

Peptide solutions were analyzed by (LC-MS/MS) on an Ultimate 3000 RSLC nano-HPLC system coupled to a Q-Exactive Plus mass spectrometer with a Nanospray Flex ion source (all from Thermo Fisher Scientific). Samples were loaded onto an RP C18 pre-column (Acclaim PepMap, 300 μm × 5 mm, 5 μm, 100 Å, Thermo Fisher Scientific) and washed with 0.1% (*v*/*v*) TFA for 15 min at 30 μL/min. Peptides were eluted from the pre-column and separated on a 200 cm µPAC C18 separation column (Pharmafluidics) that had been equilibrated with 3% solvent B (solvent A: 0.1% (*v*/*v*) FA (formic acid), solvent B: ACN, 0.1 % (*v*/*v*) FA). A linear gradient from 3–35% solvent B within 180 min at 600 nL/min (0-30 min) and 300 nL/min (30-180 min) was used to elute peptides from the separation column. Data were acquired in data-dependent MS/MS mode, HCD (higher-energy collision-induced dissociation) with a normalized collision energy (NCE) of 28% was used for fragmentation. Each high-resolution full MS scan (*m*/*z* 375 to 1799, R = 140,000, target value (TV) 3,000,000, max. injection time (IT) 50 ms) was followed by high-resolution (R = 17,500, TV 200,000, IT 120 ms, isolation window 2 Th) product ion scans, starting with the most intense signal in the full scan MS, dynamic exclusion (duration 30 s, window ± 3 ppm) was enabled.

For peptide identification and quantification, data were searched against the Swiss-Prot database (tax. *Homo sapiens,* version 11/19; 20,315 entries) using Proteome Discoverer, version 2.4 (Thermo Fisher Scientific). A maximum mass deviation of 10 ppm was applied for precursor ions, while for product ions, max. 0.02 Da were allowed. Oxidation of Met and acetylation of protein N-termini were set as variable modifications. Carbamidomethylation of cysteines was included as fixed modification. A maximum of two missed cleavage sites were considered for peptides. Peptides were quantified via LFQ (label-free quantitation) and data were finally subjected to quantile normalization to derive protein enrichment by RAVER1.

### Western Blot

Protein expression was analyzed by Western blotting as previously described (27). Primary antibodies and fluorescence-coupled secondary antibodies are indicated in supplementary table TS5.

### TGFB treatment and lethal EMT analysis

To induce an epithelial-mesenchymal-transition (EMT), cells were treated with TGF-β (TGFB). To this end, 3×10^3^ control cells or 6×10^3^ RAVER1 (higher cell numbers were required to compensate for elevated cell death and deficient proliferation) depleted cells were seeded into 96 well plate 24 h post siRNA transfection and grown for an additional 24 h. Cells were treated for 72 h with 10ng/ml recombinant human TGF-β1 (HEK293 derived, PEPROTECH). Cell density was monitored by an Incucyte S3 (Sartorius) and analyzed using the AI Cell Health Analysis Software (v2022B). Cell viability and caspase 3/7 activity was measured with CellTiter Flo (Promega) or CaspaseGlo (Promega) according to manufacturer’s protocol, respectively. Apoptosis induction was determined by normalization of caspase 3/7 activity to cell viability relative to untreated control cells to finally derive the apoptosis rate.

### Luciferase assay

Luciferase activity was analyzed 48 h post-transfection using DualGlo (Promega), essentially as previously described (25). Firefly luciferase (FFL) activities were normalized by Renilla (RL) activities yielding relative activities (RLU). An empty vector, containing only the multiple cloning site (MCS), served as negative control. RLU ratios were normalized to control populations where indicated.

### Immunostaining

Immunostaining was performed, essentially as previously described (27), using protein-specific antibodies summarized in supplementary table TS5), Phalloidin-TRITC (Sigma) and DAPI (Sigma). PNC were quantified using NuclearParticleDetector2D (MiToBo Cell Image Analyzer) within Fiji (version 1.54d) with a minimum region size of 10, essentially as previously described (28).

### RNA Sequencing and data processing

Strand-specific paired-end RNA sequencing of control, RAVER1 and PTBP1 depleted A549 cells was performed by Novogene (Hong Kong) using Illumina NovaSeq 6000, resulting in 150 bp long reads with an average of 2×28 million reads per sample.

Quality of the raw fastq files was assessed using FastQC (https://www.bioinformatics.babraham.ac.uk/projects/fastqc/). Sequencing adapters and low quality read ends were clipped off using Cutadapt v2.8. The processed sequencing reads were aligned to the human genome (UCSC hg38) using HiSat2 v2.1.0. Alignments in the obtained bam files were sorted, indexed and secondary alignments were filtered out using samtools v1.10. FeatureCounts v2.0.0 was used for summarizing gene-mapped reads. Ensembl (GRCh38.100) was used as annotation basis. Differential gene expression was determined using the R package edgeR v3.38.4 utilizing trimmed mean of M-values normalization. A false discovery rate (FDR) adjusted p-value below 0.05 was considered significant for differential gene expression. For these analyses previously reported standard pipelines were used (25).

Differential isoform expression was quantified using rMATS turbo v4.1.2 (29) using the prior generated alignment files and Gencode v29 annotations (30). Protein-coding genes were determined using the R package biomaRt v.2.56.0, as previously described (25). Sashimi plots were generated using ggsashimi v1.1.5 (31) and Gencode v29 annotations. The PSI (precent spliced in) value determines the IncLevel derived from rMATS analyses.

Hexamer frequencies within significant (½ΔPSI| > 0.05, FDR < 0.05) AS exons determined by RAVER1 and PTBP1 depletion were derived via genomic coordinates of spliced exons obtained from rMATS analyses (SE.MATS.JCEC.txt). These were extracted from determined skipped exon events in protein-coding genes identified by the R-package biomaRt, using ENSEMBL v100. Exon cDNA sequences were extracted from these coordinates using bedtools getfasta (32) and the UCSC hg38 human reference genome. cDNA sequences with lengths below the 5th and above the 95th percentile were removed and hexamer frequencies of the remaining sequences were determined via the R-package Biostrings v2.68.1. Absolute hexamer frequencies were divided by the total number of hexamers in all sequences to obtain normalized hexamer frequencies.

The enrichment of RBP binding motifs within significant AS exons were revealed by using the memes R-package utilizing the MEME suite (33). Shuffled input sequences with preserved di-nucleotide frequencies were used as background sequences. Ray2013_rbp_Homo_sapiens .dna_encoded.meme obtained from the MEME website (https://meme-suite.org/meme/) was applied as motif database.

Identification of hexamers mostly abundant in skipped exon sequences, was based on an enrichment frequency above 9e-4. Motifs were converted by the R-package Biostrings and matched against a database of RBP motifs (Ray2013_rbp_Homo_sapiens.dna_encoded.meme from https://meme-suite.org/meme/) using the runTomTom function of the memes R-package utilizing the MEME suite.

### Gene Set enrichment analyses

Gene set enrichment analyses (GSEA) was performed with the GSEA Software (version 4.2.3) and MsigDB gene sets (v2023.1) using log_2_ fold change ranked protein-coding genes from RNA-seq and mass spectrometry data with a classic scheme scoring and a minimal size of 15 or 10, respectively.

Gene ontology analyses were performed using the R package clusterProfiler v4.8.1. The enrichment of the protein-coding genes with unique splicing events (FDR < 0.05, |ΔPSI| > 0.05) in the MsigDB ontology gene set (v2023.1) was calculated and filtered for molecular functions – only enrichment with an FDR < 0.05 is shown.

### Animal handling and ethics approvals

Immunodeficient athymic nude mice (FOXN1nu/nu) were obtained from Charles River (Wilmington, MA). Animals were handled according to the guidelines of the Martin Luther University based on ARRIVE guidelines. Permission was granted by the County Administration Office for Animal Care Saxony-Anhalt (42502-2-1625 MLU). iRFP-labelled (stably transduced) parental or RAVER1 depleted A549 cells (1x 10^6^) were harvested in PBS with 50% (v/v) Matrigel (Sigma) and were injected subcutaneously (sc) into 6-week-old female nude mice (6 mice per condition). Tumor growth of isoflurane-anaesthetized mice was monitored weekly by near-infrared imaging using a Pearl imager (LI-COR, Lincoln, NE) and quantified using the Image Studio software (LI-COR). Mice were sacrificed after 25 days when parental A549 tumors reached termination criteria. Tumor volume was calculated using the formula volume = π/6 x L1 x L2 x L3, essentially as previously described (34).

### Kaplan-Meier and gene expression analyses

Hazardous ratios (HRs) for indicated genes were determined by the Kaplan-Meier (KM) plotter online tool using mRNA gene chip data for lung cancer, selected for ‘best cutoff analyses and adenocarcinoma (www.kmplot.com). Survival data of the whole TCGA tumor cohort (n=33) were performed by the Gene Expression Profiling Interactive Analysis (GEPIA) online tool (www.gepia2.cancer-pku.cn). Normalized primary tumor expression (FPKM) and associated clinical data from the TCGA project (Cancer Genome Atlas Research Network, 2017) were obtained from the GDC data portal.

### Structure Prediction and Biogrid Analyses

For prediction and visualization of the RAVER1 protein structure (E9PAU2) AlphaFold DB (35) and PyMOL Molecular Graphics System (version 2.0, Schrödinger LLC) were used. Analysis of protein association with RAVER1 (E9PAU2) was performed using bioGRID webtool version 4.4.220 (https://thebiogrid.org/).

## RESULTS

### RAVER1 promotes tumor cell proliferation

Pan-cancer analyses based on RNA-seq data provided for 33 cancers via the TCGA suggested that RAVER1 as well as PTBP1 expression is associated with unfavorable prognosis in various cancers (Figure 1A, B). Moreover, both proteins showed strongly, pan-cancer associated expression (Supplementary Figure S1A). In lung adenocarcinoma (LUAD), this trend was pronounced and the expression of both, RAVER1 and PTBP1, was significantly elevated. These findings suggested that PTBP1, potentially in concert with RAVER1, serves conserved pro-oncogenic roles. This was investigated by loss-of-function studies in a panel of carcinoma- and neuroblastoma-derived (NBL) cell lines (supplementary Table TS2). The depletion of RAVER1 revealed consistent and significant downregulation of 2D tumor cell proliferation (Figure 1C). In contrast, growth rates were only modestly, in ES-2 cells even non-significantly, affected by the depletion of PTBP1. Notably, RAVER1 knockdown slightly decreased PTBP1 protein abundance and vice versa (Figure 1C). Analyses in 3D cultured LUAD-derived A549 cells demonstrated that the depletion of RAVER1 significantly reduced spheroid growth and anoikis resistance, suggesting that RAVER1 is a conserved enhancer of tumor cell proliferation (Figure 1D, E). Whether RAVER1 or PTBP1 influence cell cycle progression was evaluated in three tumor cell lines (A549, ES-2 and HepG2). RAVER1 depletion led to significant enrichment of all cell lines in the G1 phase (Figure 1F; Supplementary Figure S1B). Moreover, RAVER1 downregulation elevated caspase 3/7 activity, suggesting induction of apoptosis (Figure 1G). Taken together, this supported a conserved, pro-proliferative and anti-apoptotic role of RAVER1 in cancer cells, which appeared less pronounced for PTBP1.

**Figure 1.**
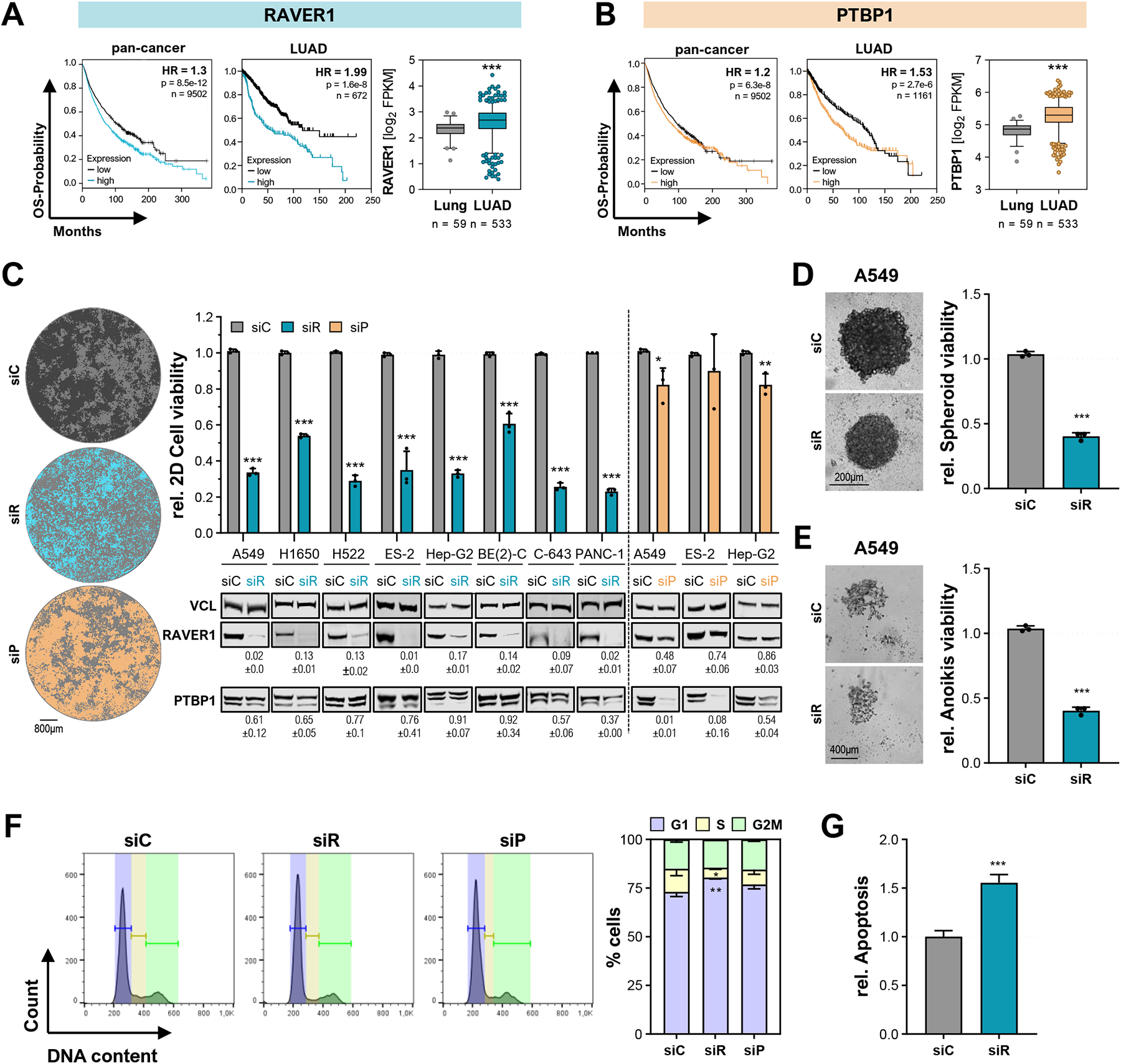
RAVER1 promotes tumor cell proliferation. **(A, B)** Overall survival Kaplan-Meier and box plots of RAVER1 (A) and PTBP1 (B) mRNA expression in 33 cancers (left panels) and LUAD (middle and right panels). HR, Hazard ratio; p, log rank p value; n, number of analyzed samples. **(C)** Proliferation analyses by CellTiter GLO (upper panel) in indicated cancer cell lines upon control (siC, gray), RAVER1-directed (siR, turquoise), and PTBP1-directed siRNA pools (siP, apricot). Representative A549 2D cell confluences 6 days post-transfection is indicated in the left panel. Black dots represent median-normalized values of three independent replicates. Representative Western blot analyses of RAVER1 and PTBP1 upon depletion (72h) are shown in the bottom panel. Vinculin (VCL) serves as loading and normalization control to derive the fold downregulation of indicated proteins in three independent analyses. **(D, E)** Spheroid growth (D) and anoikis-resistance (E) of RAVER1-depleted A549 was determined by CellTiter GLO. Representative images are shown in the left panels. **(F)** Cell cycle progression analyses of control-, RAVER1- or PTBP1-depleted A549 cells. Fractions of A549 cells in indicated cell cycle phase were quantified in three independent studies. **(G)** Induction of apoptosis upon RAVER1 depletion in A549 was determined by caspase 3/7-activity. Black dots represent median-normalized values of three independent replicates. Error bars indicate standard deviation derived from three independent analyses. Statistical significance was determined by Student’s t-test: (*) P < 0.05, (**) P< 0.01, (***) P < 0.001.

### RAVER1 deletion impairs tumor growth

To evaluate the oncogenic potential of RAVER1 in tumor growth, we established RAVER1-deleted A549 cell clones which notably showed barely affected PTBP1 expression (Figure 2A). Consistent with decreased proliferation observed upon transient RAVER1 depletion by siRNA pools, RAVER1 knockout (KO) decreased 2D proliferation as well as spheroid growth and most prominently anoikis resistance (Figure 2B-D). How RAVER1 influences tumor growth was investigated by subcutaneous (s.c.) xenograft models in nude mice (Figure 2E). To visualize tumor growth by non-invasive imaging, parental and RAVER1-KO A549 cells were stably transduced with iRFP (near infrared fluorescent protein) before s.c. injection (Figure 2F). The assessment of tumor growth by volume as well as endpoint tumor mass (after 25d) indicated that RAVER1-KO led to significantly reduced tumor volume already two weeks after injection (Figure 2G, H). Tumors derived from parental A549 cells reached termination criteria at ∼25d post-injection, whereas volume remained ∼2-fold smaller for RAVER1-KO derived tumors. Consistently, final tumor mass was reduced (p = 0.07) by RAVER1-KO, providing strong evidence that RAVER1 serves pro-oncogenic roles in LUAD xenograft models by enhancing or maintaining elevated proliferation and tumor growth.

**Figure 2.**
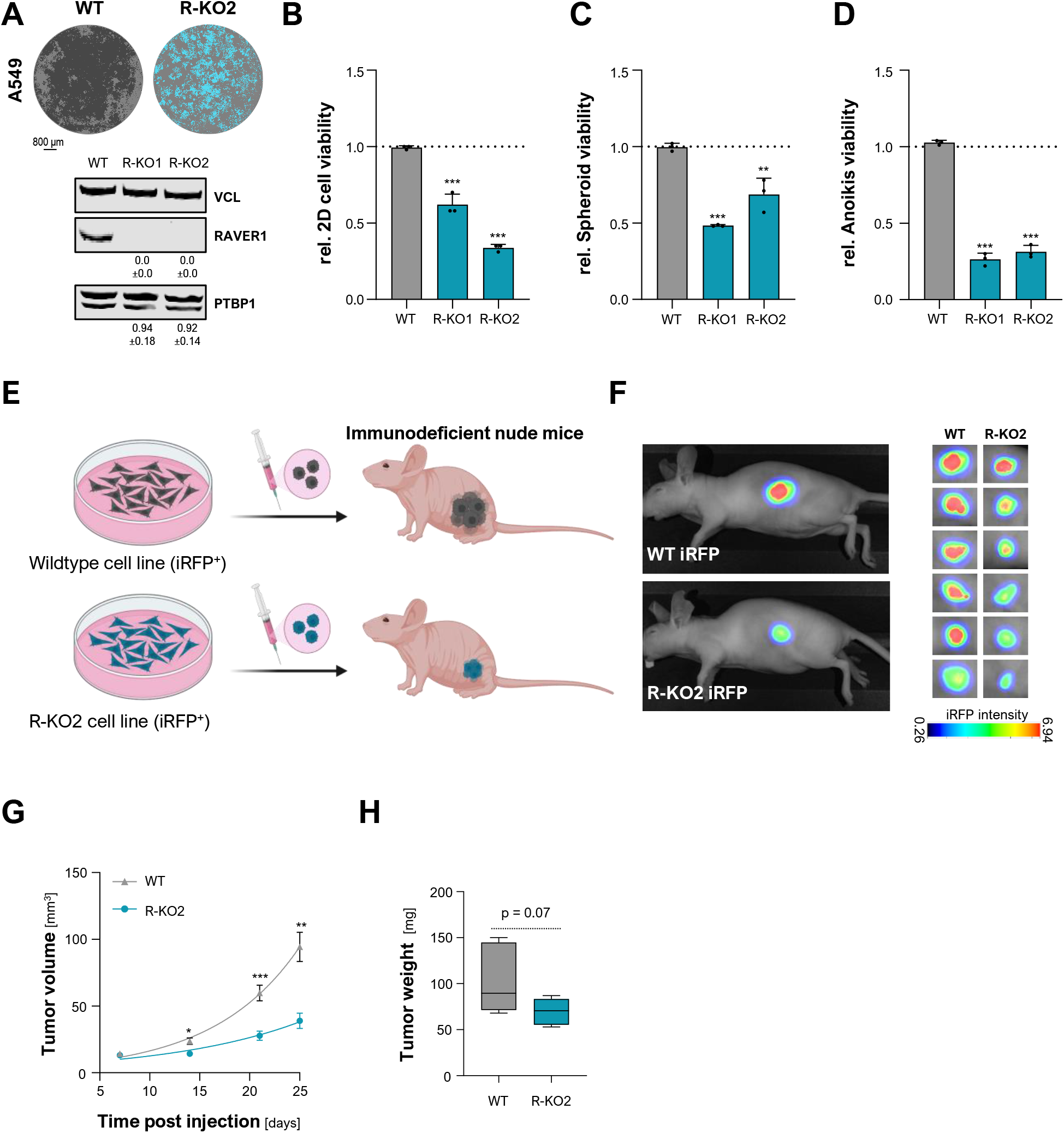
RAVER1 deletion impairs tumor growth. **(A)** Representative cell confluences (upper panel) and Western blot analysis (lower panel) demonstrating deletion of RAVER1 by CRISPR/Cas9 in two independent A549 cell clones (R-KO, turquoise) versus parental cells (WT, black). Quantification of Western blots as described in Figure 1C. **(B-D)** 2D cell proliferation (B), spheroid growth (C), and anoikis-resistance (D) were determined in RAVER1-KO (R-KO, turquoise) and parental cells (WT, gray) as in Figure 1C. **(E-H)** Parental (WT, grey) and RAVER1-deleted (R-KO2, turquoise) A549 cells expressing iRFP were injected (sc) into nude mice (6 per condition), as schematically shown in (E). Scheme was created with BioRender (ZP25HKUDFM). Tumor growth of xenograft was monitored by near-infrared imaging. Representative images 25 days post injection are shown in (F). Tumor volumes over time (G) and final tumor mass in (H) are shown. Error bars indicate standard deviation derived from three independent analyses. Statistical significance was determined by Student’s t-test: (*) P < 0.05, (**) P< 0.01, (***) P < 0.001.

### RAVER1 is dispensable for perinucleolar compartment (PNC) homeostasis

The original identification of RAVER1 revealed its association and co-localization with PTBPs in perinucleolar compartments (17). Independent studies suggested elevated frequency of such PNCs in progressed malignancies and associated with enhanced metastatic potential in cancer (22,23). Accordingly, it was tempting to speculate that RAVER1’s role in tumor cell proliferation is associated with its potentially PTBP-dependent modulation of PNC homeostasis. Initially, this was addressed by revisiting RAVER1-PTBP1 association in RAVER1-KO A549 cells re-expressing either wild-type or mutant RAVER1 proteins (Figure 3A). Affinity purification of SBP-FLAG tagged RAVER1 proteins from nuclear fractions showed that the wild-type protein and a mutant lacking the RRMs associated with PTBP1 as expected (Figure 3B). Deletion of RAVER1’s intrinsically disordered region (IDR) as well as mutation of prior described PTBP1-interacting motifs within the IDR abrogated this association, although previously identified NLS and NES motives were maintained.

**Figure 3.**
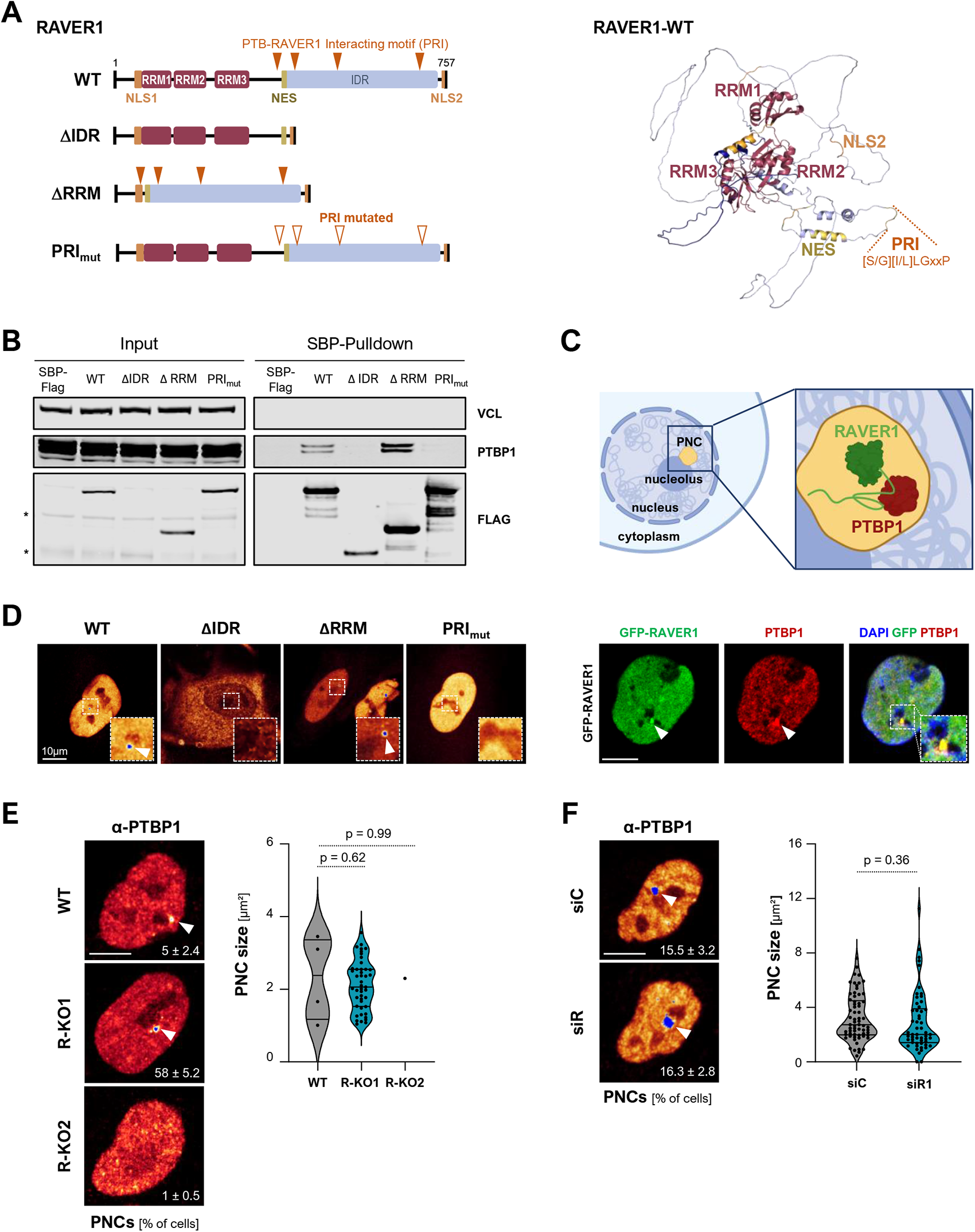
RAVER1 is dispensable for PNC homeostasis. **(A)** Alphafold structure prediction of RAVER1 (E9PAU2, right) and scheme of RAVER1 proteins (left). RRM, RNA-Recognition Motif; NLS, Nuclear Localization Site; NES, Nuclear Export Signal. **(B)** Representative Western blot analysis of SBP-Immunoprecipitation from nuclear extracts of A549 R-KO cells re-expressing RAVER1 proteins indicated in (A). (*)unspecific background staining. **(C)** Localization studies of GFP-RAVER1 (stably expressed in A549 R-KO) and PTBP1 by immunostaining (lower panel) and schematic of RAVER1-PTBP1 association in PNCs (upper panel, created with BioRender: YT25HKUPUH). **(D)** Immunostaining (α-SBP) of RAVER1 proteins shown in (A) and re-expressed in A549 R-KO cells. Dashed boxes depict enlarged region shown in the upper right corner. **(E, F)** Immunostaining of PTBP1 in parental (WT) and RAVER1-deleted (R-KO) A549 cells (E, n=80), and upon control (siC) and RAVER1-depletion (siR) in A549 cells (F, n=300). Violin plots depict PNC size distribution determined by NuclearParticleDetector (Fiji). Scale bar (C-F) 10 µm. Statistical significance was determined by Student’s t-test.

PTBPs and RAVER1 are co-localized at PNCs in A549 cells (Figure 3C), as previously demonstrated in other cell models (17). This was also observed for the SBP-FLAG-tagged RAVER1 wild-type protein re-expressed in RAVER1-KO A549 cells (Figure 3D). Strikingly, only RAVER1 proteins containing the intact IDR showed localization in PNC-like nuclear structures. This supported the notion that the IDR-dependent association of RAVER1 with PTBPs is instrumental for the PNC-localization of RAVER1, PTBP1 and potentially PNC integrity. Whether RAVER1 is essential for PNC formation and PTBP1-recrutiment was analyzed by RAVER1-KO and knockdown in A549 cells (Figure 3E, F). Immunostaining for PTBP1 in parental as well as RAVER1-KO cells revealed a strong clonal variability of PNC frequency, although both clones showed similarly decreased proliferation compared to parental cells (cf. Fig.2B-D). In parental populations only ∼5% of cells contained PTBP1-positive PNCs, whereas nearly 60% of cells showed PTBP1-containing PNCs in RAVER1-KO#1 but only ∼1% of PNC-positive cells was observed in RAVER1-KO#2. However, despite variable frequency of PNCs, their size distribution was essentially unchanged. This was tested further by RAVER1 depletion using siRNA pools. However, PTBP1-recrutiment to PNCs as well as PNC frequency and size distribution remained essentially unaffected by RAVER1 knockdown. This demonstrated that RAVER1 is dispensable for PNC integrity, PNC-recruitment of PTBP1 and controls tumor cell proliferation in a largely PNC-independent manner.

### RAVER1 depletion impairs proliferation pathways and promotes lethal EMT

RAVER1 depletion and knockout disturbed tumor cell proliferation and tumor growth. In agreement, gene set enrichment analyses (GSEA) derived by RNAseq data in RAVER1-depleted A549 cells confirmed severe downregulation of pro-proliferative gene expression upon knockdown (Figure 4A; Supplementary Table TS6 and TS7). Essentially all pro-proliferative cancer hallmark gene sets (MsigDB-HM), most prominently E2F_target_genes (HM-E2F), were significantly decreased by RAVER1 knockdown, supporting the prior identified disturbance of G1/S transition and cell cycle progression (cf. Fig.1). Surprisingly, however, GSEA suggested induction of epithelial-mesenchymal-transition (EMT; HM_EMT) presumably due to elevated TGFB-signaling (HM_TGFβ) upon RAVER1 as well as PTBP1 knockdown. This was supported by the upregulation of mesenchymal markers like CDH2 (N-cadherin) and SNAI2 (Slug), whereas classical epithelial markers like CDH1 (E-cadherin) were substantially decreased (Figure 4B; Supplementary Table TS6). The upregulation of both, TGFBRI and TGFBRII, suggested auto- or paracrine induction of TGFB-driven EMT. For PTBP1 knockdown, the analysis of cell morphology, F-actin remodeling, the subcellular localization of CDH2 and the multi-versatile adherens junction protein CTNNB1 (β-catenin) demonstrated no substantial changes (Figure 4B; Supplementary Figure S2A). In contrast, RAVER1 depletion enhanced a mesenchymal-like cell morphology. This was evidenced by strong enrichment of CDH2 and loss of CTNNB1 at the cell periphery in favor of elevated cytoplasmic localization. Concomitantly, the actin cytoskeleton was shifted from epithelial-like enrichment of rather thin peripheral F-actin structures to mesenchymal-like, strong stress-fibers traversing the cytoplasm. Collectively, these findings provide strong evidence that RAVER1 depletion promoted EMT, whereas PTBP1 affected the same overall pathways but apparently is less essential in maintaining epithelial integrity. However, EMT is a hallmark of carcinoma progression, yet tumor cells proliferated slower, and apoptosis appeared elevated upon RAVER1 downregulation whereas this was less pronounced by PTBP1 knockdown (cf. Fig.1). This suggested that RAVER1 depletion, in contrast to PTBP1, shifted TGFB signaling strongly towards the induction of apoptosis, which would be concise with the induction of lethal EMT. In support of this, GSEA analyses in the MsigDB-C2 collection revealed a variety of TGFB-associated gene panels, which showed and overall similar enrichment (Pearson correlation: R = 0.6043, p < 0.001) for both, RAVER1 as well as PTBP1 depletion (Figure 4C). However, two gene sets, both comprising changes in gene expression reported as lethal EMT or TGFB-induced apoptosis (36), stood out due to enrichment by RAVER1 depletion, whereas this was not seen for PTBP1 knockdown. Consistent with the original reports (36), RAVER1 depletion induced a strong enhancement of TGFB-driven facilitators of apoptosis, BIM (BCL2L11) and BMF (Figure 4C). In addition, gene expression changes supported another, PDAC-derived (pancreatic ductal adenocarcinoma) report on lethal EMT induction by TGFB (37). LUAD-derived A549 cells are KRAS-mutated (G12S), a disturbance frequently observed in PDAC, and maintain SMAD4, SOX4 as well as KLF5 synthesis (Figure 4C). RAVER1 depletion upregulated TGFB signaling, SNAI2, and SOX4 but repressed KLF5 (Figure 4B, C), which was reported to impose the induction of apoptosis (37). Elevated TGFB signaling and EMT induction likely resulted from enhanced expression of TGBFRI/II and SNAI2, two essential drivers of cancerous EMT (38). If RAVER1 knockdown promoted TGFB-induced, lethal EMT was further tested in A549 cells exposed to TGFB by monitoring caspase 3/7 activity (apoptosis) and cell viability (Figure 4D). In controls, TGFB treatment barely affected cell viability or apoptosis. RAVER1 depletion, however, substantially elevated apoptosis rates upon TGFB exposure, demonstrating that RAVER1 antagonizes lethal EMT, presumably by restricting auto- and paracrine TGFB/BMP signaling and the induction of lethal EMT (Figure 4E).

**Figure 4.**
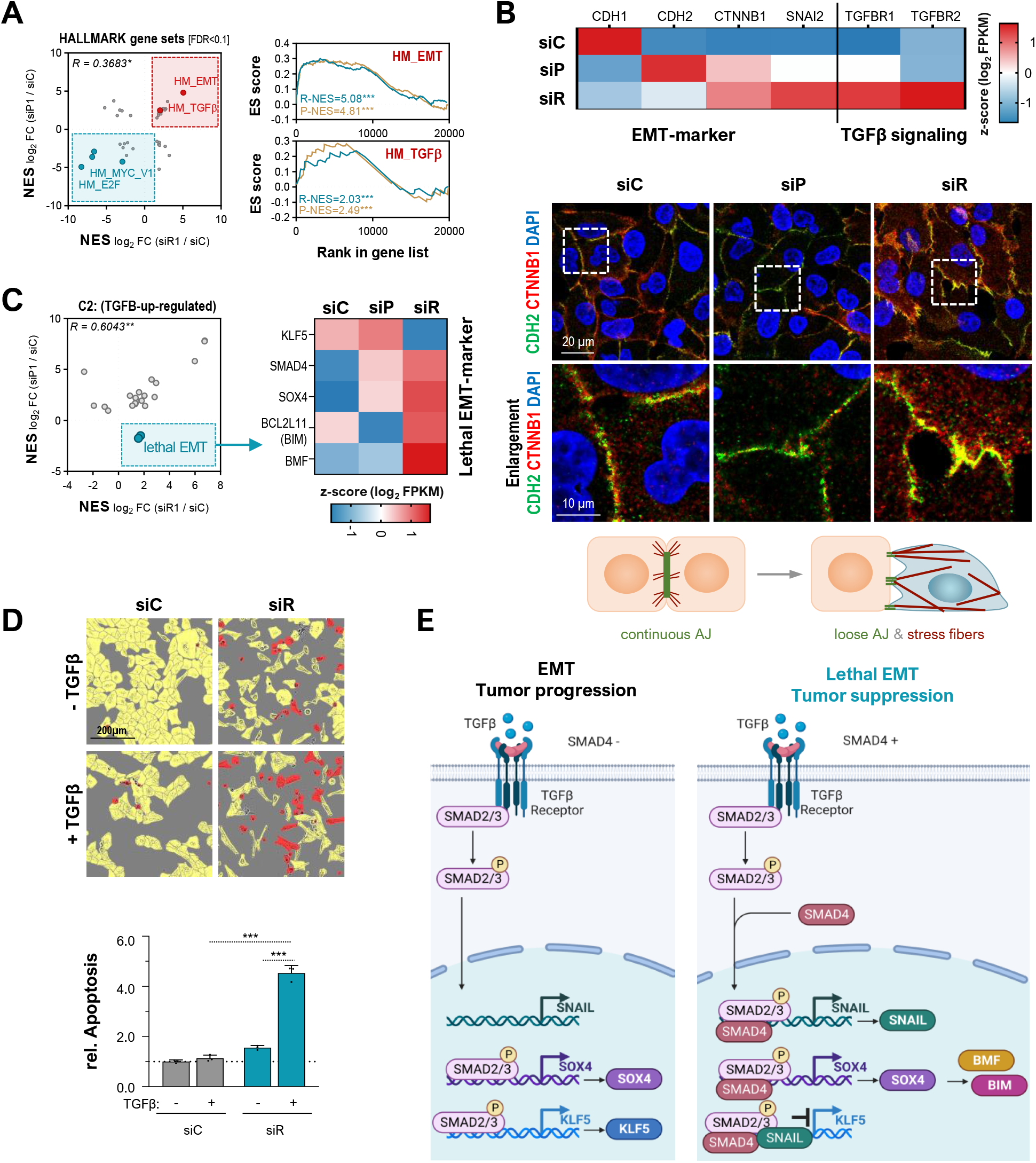
RAVER1 depletion impairs proliferation pathways and promotes lethal EMT. **(A)** XY-presentation of NES values for Hallmark (HM) gene sets determined by GSEA upon RAVER1 (turquoise) and PTBP1 depletion (apricot) in A549 cells. **(B)** Analysis of selected EMT-Marker by RNA-seq (upper panel) and immunostaining of CHD2 and CTNNB1 upon RAVER1- or PTBP1 depletion in A549 cells. Gene expression changes are indicated as median z score (log2 FPKM) in the upper panel. A schematic depicting altered adherens junction (AJ) morphology is in the lower panel. **(C)** XY-presentation of NES scores for TGFB-associated gene sets derived by GSEA in the C2 collection as described in (A). **(D)** Apoptosis induction (lower panel) upon TGFβ treatment in control (gray) and RAVER1 depleted (turquois) A549 cells. Representative confluence images (upper panel) of living (yellow) and apoptotic (red) cells, as determined by IncuCyte AI quantification. **(E)** Scheme of TGFβ-induced cancerous EMT (left) and lethal EMT (right), as proposed by (36,37). Scheme was created with BioRender (SY25HKUS7O). Error bars indicate standard deviation derived from three independent analyses. Statistical significance was determined by Student’s t-test: (*) P < 0.05, (**) P< 0.01, (***) P < 0.001.

### RAVER1 modulates activity of the miR/RISC-guided control of gene expression

Aiming to unravel how RAVER1 and PTBP1 influence TGFB signaling, although to distinct extend, we analyzed gene set enrichment in the MsigDB-C3 collection, containing gene panels with shared transcription factor targeting (TFT) as well as microRNA (miR, miRNA) targeting motifs. Strikingly, for miRNAs expressed in A549 cells (cpm > 10), RAVER1 as well as PTBP1 knockdown increased essentially all target gene panels (Figure 5A; Supplementary Figure TS7). In contrast TFT gene panels showed a rather balanced distribution but overall downregulation. Inspection of the top five most depleted and enriched gene panels, however, emphasized substantial differences for RAVER1 versus PTBP1 knockdown (Figure 5B). In contrast to PTBP1, the downregulation of RAVER1 depleted various E2F-target gene sets among the five most depleted gene panels, supporting a pronounced role in cell cycle progression, in particular G1/S transition (cf. Fig.1). Among enriched gene sets, PTBP1 as well as RAVER1 knockdown elevated miR-target gene panels, supporting previous findings that suggest PTBP1 as a modulator of miR/RISC-guided mRNA turnover (39). However, in contrast to PTBP1, RAVER1 knockdown most prominently increased targets of the miR-200 family and the oncomiR-21. The miR-200 family is central for TGFB-induced cancerous EMT and maintaining epithelial integrity (40,41). This suggested that RAVER1 influences TGFB-signaling and EMT largely by modulating miR/RISC activity in a potentially mRNA dependent manner. Accordingly, RAVER1 knockdown either led to disturbed miR biogenesis, turnover, or deregulated miR/RISC activity in gene silencing. Northern blotting of total RNA isolated from RAVER1 and PTBP1 knockdown cells as well as RAVER1-KO clones showed that the abundance of miR-200b-3p, let7a-5p as well as miR-21-5p, the most abundant microRNA in carcinomas (42), remained essentially unchanged (Figure 5C). Thus, it appeared unlikely that the extensive upregulation of miR-target transcripts in knockdown cells resulted from broad downregulation of miRNA abundance. How RAVER1 knockdown affects miR/RISC activity was initially tested by minimal luciferase reporters harboring three, tailed, perfectly complementary miR-21-5p or let-7a targeting sites in their 3’UTR (Figure 5D). RAVER1 knockdown significantly, but moderately increased the activity of these reporters. The depletion of RAVER1 also elevated the activity of reporters comprising entire 3’UTRs (Figure 5D). These were derived from major miR-controlled mRNAs like HMGA2 and LIN28B, mainly regulated by the let-7 family (24), and CD274/PD-L1 as well as MDM2, regulated by multiple miRs (43,44). Consistent with upregulated reporter activity the corresponding mRNAs were enhanced by RAVER1 depletion next to various other mRNAs, with major, reported miR-dependent regulation: IGF2BP1 (mainly let-7, (24)), ZEB2 and TGFBR2 (mainly miR-200 family, (40,41)), tumor-suppressive BTGs (mainly miR-21 (42)) (Figure 5E). Notably, however, we also observed that major miR-target mRNAs, e.g. PDCD4 (miR-21, (45)), were decreased by RAVER1 knockdown, suggesting mRNA-dependent secondary regulation. Despite these divergent observations, however, our findings provided strong evidence that RAVER1 and potentially PTBP1 modulate miR/RISC-activity in a partially mRNA-dependent manner.

**Figure 5.**
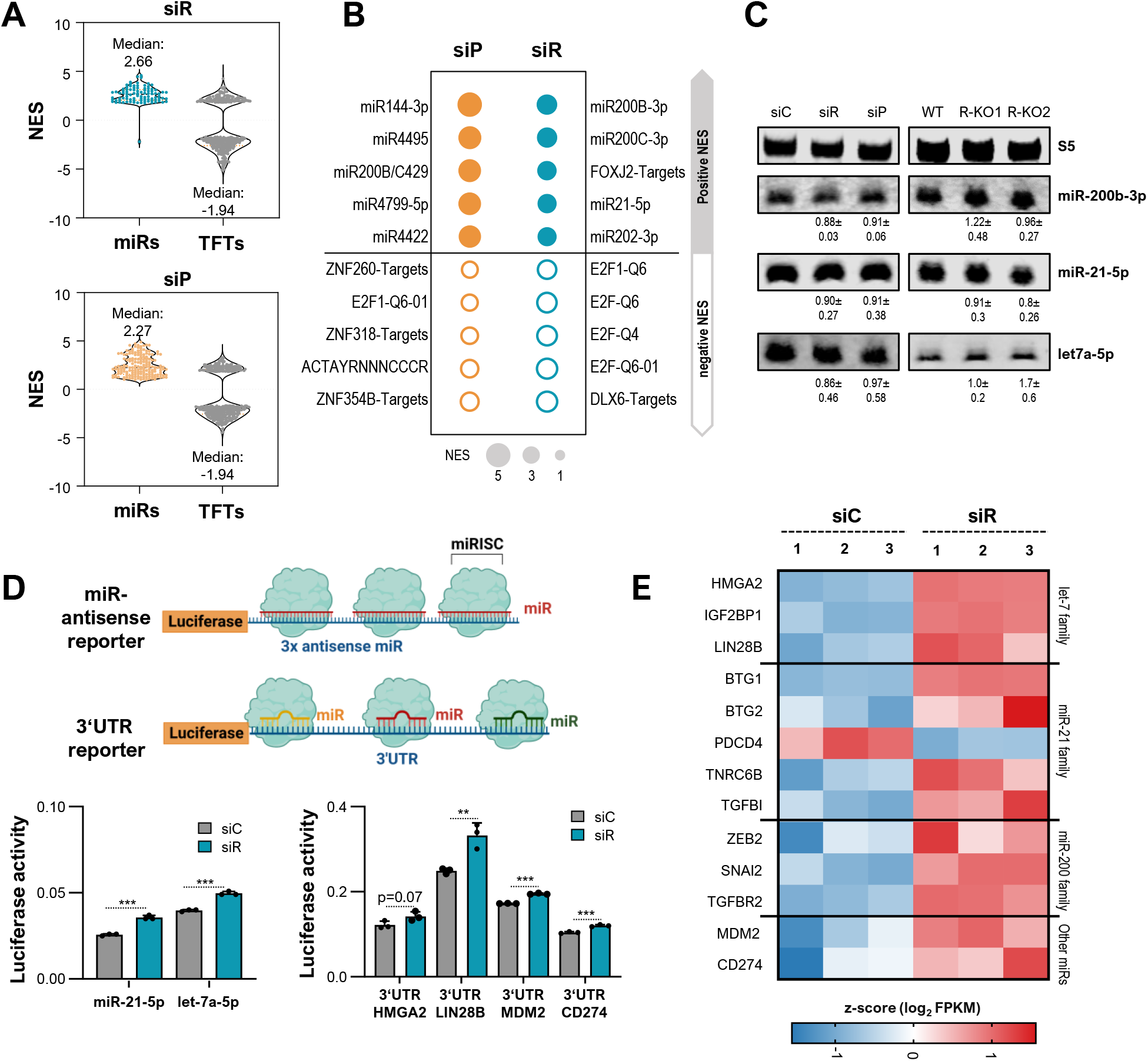
RAVER1 modulates the miR/RISC-guided control of gene expression. **(A)** Violin Plots of NES values determined by GSEA in the C3-MIR (colored) and C3-TFT (gray) collections upon RAVER1 (upper panel) and PTBP (lower panel) depletion in A549 cells. Only NES scores with FDR<0.05 were considered. **(B)** Top five C3-all gene sets by NES score as derived by GSEA upon PTBP1 (left) and RAVER1-(right) depletion in A549 cells. **(C)** Northern blot analysis of indicated microRNAs upon RAVER1- and PTBP1-depletion (left), as well as RAVER1-deleltion (right) in A549 cells. Average fold changes and standard deviation of microRNA levels determined in three independent analyses are indicated. **(D)** Activity of indicated luciferase reporter upon control (gray) and RAVER1 depletion (turquoise) in A549 cells (n=3) in the lower panel. Schemes (created with BioRender: KV25HKUK72) depicting used reports are presented in the upper panel. Reporter activities were normalized to control reporter without targeting site (not shown). **(E)** Heat map of altered abundance of indicated mRNAs as described in Figure 4B. Error bars show standard deviation derived from three independent analyses. Statistical significance was determined by Student’s t-test: (**) P< 0.01, (***) P < 0.001.

### RAVER1 modulates alternative splicing in synergy with and independent of PTBP1

The presented findings suggested that RAVER1 is a key regulator of the miR/RISC pathway. However, the investigation of transcripts encoding effectors of miR/RISC-directed regulation did not reveal any obvious changes in abundance supporting broad impairment of miR/RISC activity observed upon RAVER1 knockdown (Supplementary Figure 3A). Accordingly, it was tempting to speculate that RAVER1 depletion disturbed miR/RISC activity by shifting alternative splicing (AS). Therefore, splicing was investigated by RNAseq in A549 cells upon RAVER1 as well as PTBP1 knockdown and data evaluation by rMATS (29).

Consistent with shRNA-directed PTBP1 knockdown in liver cancer-derived Hep-G2 cells (46), transient PTBP1 depletion by siRNAs in A549 cells predominantly impaired exon skipping and inclusion (Figure 6A; Supplementary Table TS8). This was also observed for RAVER1 knockdown, providing the first comprehensive evidence that RAVER1 mainly functions in the modulation of AS. The investigation of significant AS events (FDR < 0.05, |ΔPSI| > 0.05) in protein-coding genes revealed that both, RAVER1 as well as PTBP1 knockdown, promoted skipping as well as inclusion events in a nearly balanced manner and essentially indistinguishable exon width distribution (Figure 6B, C; Supplementary Figure S3B, C).

**Figure 6.**
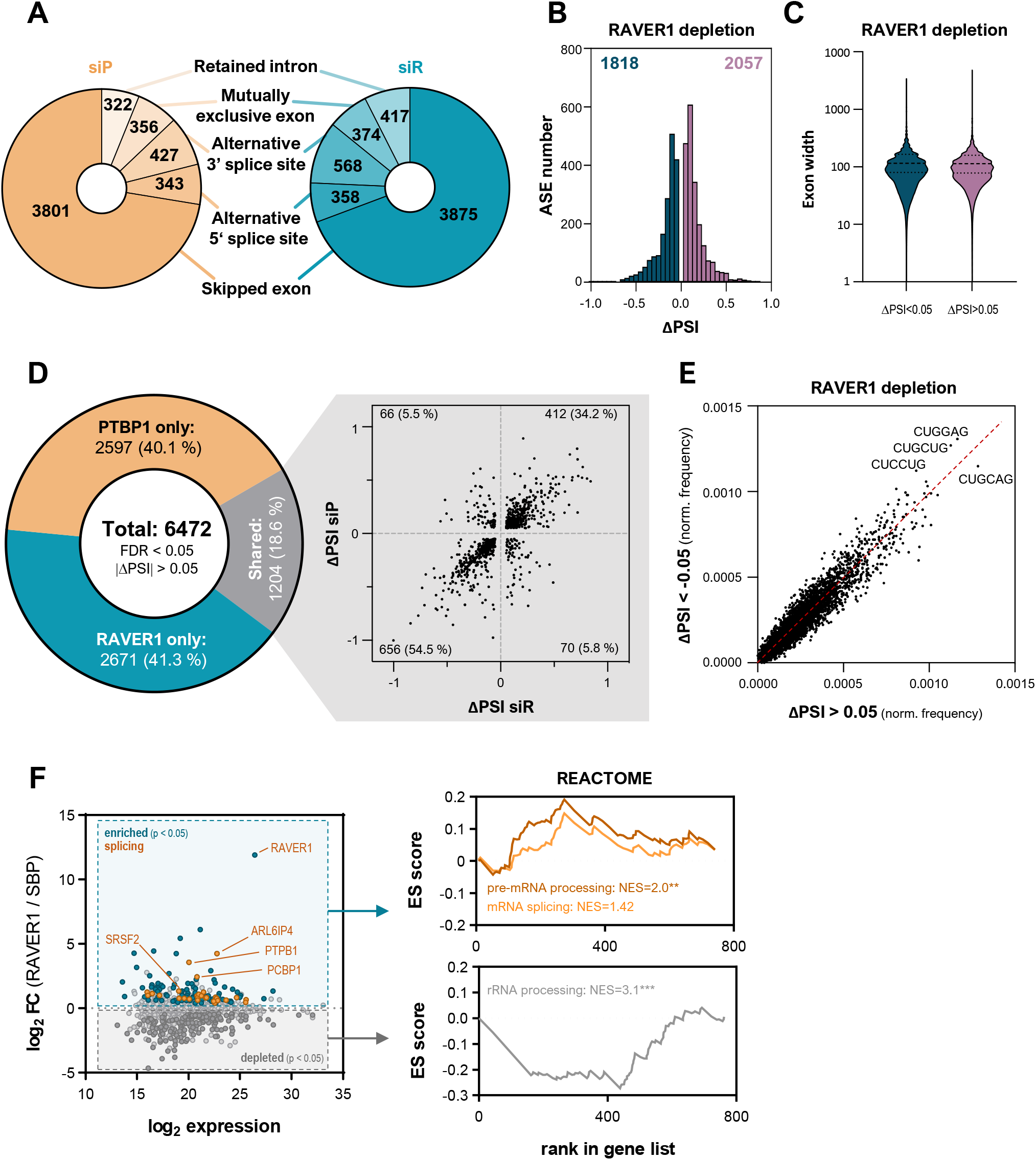
RAVER1 modulates alternative splicing. **(A-C)** Analysis of significant rMATS-reported splicing events upon RAVER1-(turquoise) and PTBP1-(apricot) depletion in A549 cells. Events were filtered for protein coding transcripts, FDR < 0.05 and |ΔPSI| > 0.05. Distribution of indicated splicing events are shown in (A). Plots depicting the frequency of alternative splicing (AS) events by ΔPSI values (B) and exon width (C) upon RAVER1 depletion are shown. **(D)** Distribution of indicated AS events with ΔPSI correlation plot for shared events upon RAVER1 and PTBP1 depletion. **(E)** Normalized hexamer frequency in exons for indicated ΔPSI AS events in XY-presentation. **(F)** LC-MS/MS analyses of SBP-RAVER1 immunoprecipitation (IP) was used to identify RAVER1 protein interaction partners. Enrichment of RAVER1-IP over control is shown as MA-plot and significantly altered proteins (p < 0.05) as well as splicing regulators are indicated by color (left panel). GSEA using C2/Reactome gene sets identified “Reactome processing of capped intron containing pre-mRNA” (orange) and “rRNA processing” (gray) among the most altered gene sets.

The comparison of 6472 significant (FDR 0.05, |ΔPSI| > 0.05) AS events in protein coding genes observed by RAVER1 and PTBP1 depletion unraveled an unexpectedly small overlap of 1204 (18.6 %) AS events (Figure 6D). These, however, showed an expected coherent regulation, with strong Pearson correlation (R = 0.72, p < 0.0001) and more skipping (656, 54.5%) than inclusion (412, 34.2%), supporting previously reported co-regulation in AS by both proteins. GSEA analyses based on gene ranking by PSI values failed to reveal any enrichment of gene panels among coherently, shared AS events (data not shown). GO term analyses of PTBP1_only, RAVER1_only or shared target genes suggested a moderate enrichment of genes encoding proteins implicated in cytoskeletal protein binding among shared and PTBP1_only events. This was slightly distinct for RAVER1 which preferentially modulated factors involved in catalytic_acting_on_DNA activity (Supplementary Figure S3E and Table TS9). Thus, despite some overlapping regulatory roles these findings implied that RAVER1 and PTBP1 mainly serve independent roles in the control of AS.

Hexamer motif analyses in exons differentially regulated by RAVER1 and PTBP1 indicated partially distinct motif enrichment in skipping versus inclusion events (Figure 6E; Supplementary Figure S3D and Table TS10). The range of enrichment was similar to hexamer distribution observed for SRFS4/6-dependent, hypoxia-associated AS in cancer cells (47). Strikingly, CU-enriched motifs were enhanced within AS exons deregulated by RAVER1 as well as PTBP1 knockdown. Likewise, a similar, yet in part distinct enrichment was observed for RBP-binding motifs within AS exons, especially motifs reported for other splicing regulators like SRSFs (Supplementary Table TS10). These findings suggested that RAVER1, like proposed for its association with PTBP1 (18), modulates splicing by complexing with a variety of other splicing regulators. According to protein association reported by BioGRID, RAVER1 was proposed to associate with various RBPs, including SRSFs (Supplementary Table TS11). This was analyzed in further detail in RAVER1-KO A549 cells, re-expressing SBP/FLAG-tagged RAVER1 or SBP/Flag, serving as control. The investigation of proteins selectively co-purified with RAVER1 from nuclear fractions by LC-MS/MS confirmed purification of RAVER1 and association with PTBP1 (Figure 6F; Supplementary Figure 3F). In addition, RAVER1 bound SRFSs, for instance SRSF2 which showed enriched motifs in RAVER1-controlled exons (Supplementary Table TS10 and TS12), and other splicing regulators like PCBP1. In agreement, GSEA in the C2-reactome collection indicated enrichment of mRNA processing and splicing regulators among RAVER1 associated proteins (Figure 6F; Supplementary Table TS12).

### RAVER1 is a master regulator of alternative splicing within the miR/RISC pathway

The analysis of gene enrichment among deregulated splicing events upon RAVER1 and PTBP1 depletion did not highlight factors implicated in the control of miR/RISC activity. Therefore, we investigated a list of 29 factors with validated roles in miR biogenesis, miR/RISC activity, and miR decay. Strikingly, hypergeometric testing indicated a ∼3.3-fold enrichment (20/29 genes, p = 4e^-08^) of disturbed AS within the miR/RISC pathway upon RAVER1 depletion (Figure 7A, B; Supplementary Table TS13). This deregulation was less pronounced for PTBP1 knockdown (∼1.9-fold enrichment, 11/29 genes, p = 0.02). The evaluation of selected AS events by semi-quantitative RT-PCR largely confirmed disturbed isoform synthesis, as determined by rMATS (Figure 7C, D; Supplementary Figure S4A). In addition, to changing relative isoform abundance, RAVER1 knockdown also altered the overall abundance of factors modulating miR/RISC activity. This was most prominently observed by significant upregulation of all three TNRC6 proteins (Supplementary Figure S3A and Table TS6). Notably, most splicing events within miR/RISC pathway genes were mainly (e.g. TNRC6B) or exclusively (e.g. TNRC6C) controlled by RAVER1, although we also observed co-regulation by RAVER1 and PTBP1 for some factor, e.g. TNRC6A.

**Figure 7.**
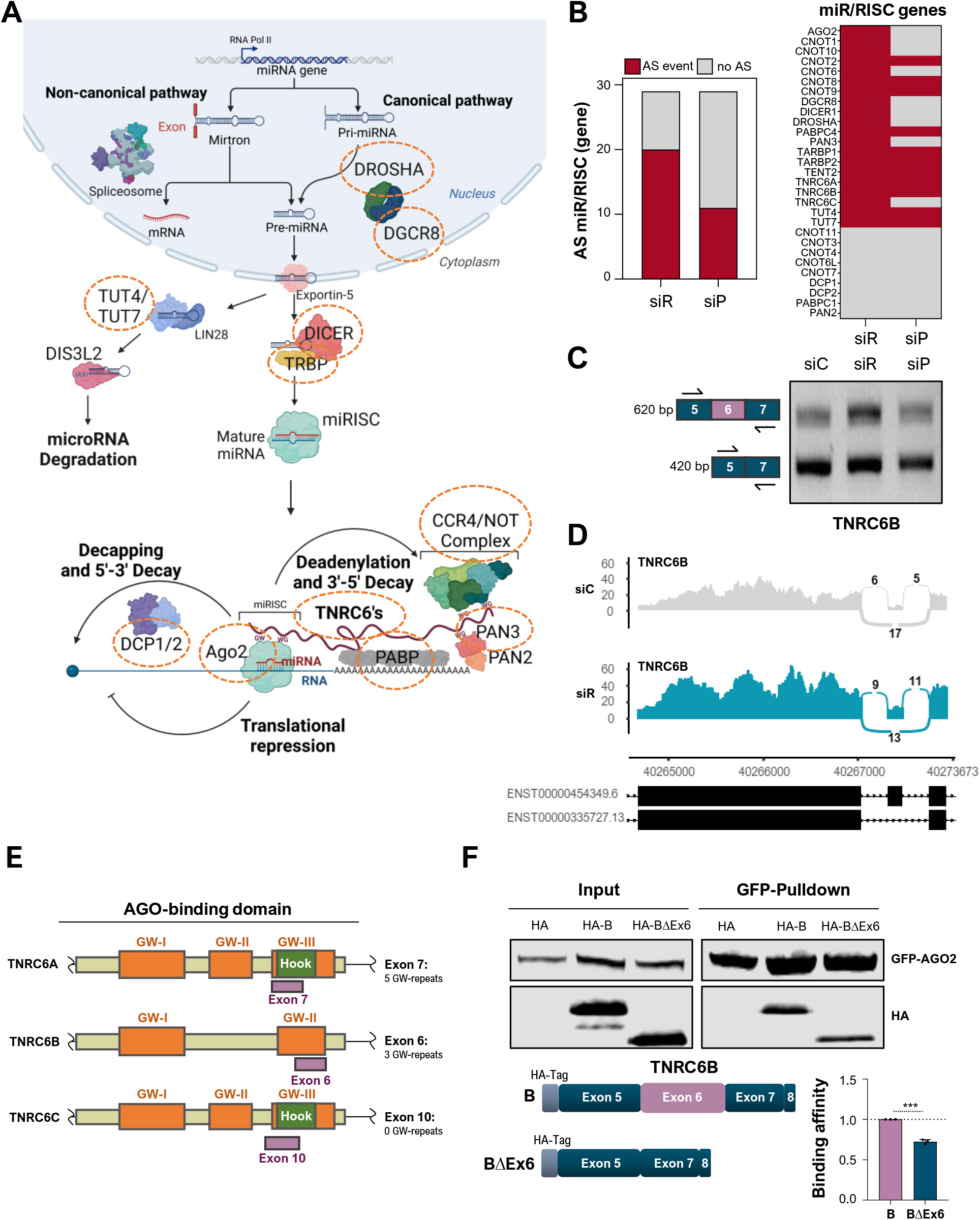
RAVER1 is a master regulator of alternative splicing within the miR/RISC pathway. **(A)** Scheme of key regulators in miR/RISC-directed gene silencing, created with BioRender (YL25HKU4MT). Orange circles indicated factors with significant AS events upon RAVER1 depletion in A549 cells. **(B)** Bar plot and heat map (right) indicating the number and miR/RISC pathway genes with significant AS events (red) upon RAVER1- and PTBP1-depletion in A549 cells. **(C)** PCR analysis of exon 6 splicing in TNRC6B (scheme in left panel) in response to RAVER1-(turquoise) and PTBP1-(apricot) depletion in A549 cells. **(D)** Sashimi plot of TNRC6B splicing in RAVER1 depleted A549 cells, as determined by rMATS. **(E)** Scheme of protein GW-rich regions in TNRC6 proteins with alternatively spliced exons. **(F)** Co-immunoprecipitation of GFP-AGO2 with indicated (scheme in lower, left panel) truncated, HA-tagged TNRC6B (B: TNRC6B, BΔEx6: TNRC6B without Exon 6), proteins. Representative Western blot analysis (upper panel) and quantification of TNRC6B copurification determined in three independent studies are shown in a bar diagram (lower, right panel). Error bars show standard deviation derived from three independent analyses. Statistical significance was determined by Student’s t-test: (***) P < 0.001.

The most prominent and concise control of exons among genes implicated in guiding miR/RISC activity was determined for TNRC6 proteins. These serve largely redundant roles in modulating miR/RISC activity by orchestrating the link between mRNA-associated miRISC complexes and the 3’end decay machinery (Figure 7A, (48,49)). Essential for this bridging function of TNRC6 proteins is their association with AGO2 via GW-enriched, intrinsically disordered regions, present within all three TNRC6 paralogues (Figure 7E). The downregulation of RAVER1 modulated alternative splicing of AGO2-binding GW-rich motifs (GW-II or GW-III, also termed AGO hook domain) in all three TNRC6 proteins. If alternative composition of the AGO hook domain indeed affects AGO2 association was analyzed for exon 6 of TNRC6B. IP studies from co-transfected HEK293 cells showed that deletion of this exon significantly reduced GFP-AGO2 association of truncated TNRC6B proteins, comprising GW-II (Figure 7F). Collectively, these studies suggested that RAVER1 modulates miR/RISC activity by guiding alternative splicing of various miR/RISC pathway genes, specifically factors involved in gene silencing by miR/RISC complexes.

## Discussion

The RAVER1 protein has been considered one of several co-factors synergizing with the key splicing regulatory PTBP protein family (16,17), which has been implicated in cancer progression (10). In this study, we provide the first evidence that RAVER1 is a pro-oncogenic modulator of alternative splicing (AS) in cancer cells, which controls splicing in concert as well as independent of PTBP1. Unexpectedly, RAVER1 depletion/deletion impaired tumor cell fitness in a more robust and conserved manner than PTBP1 knockdown. In s.c. LUAD (A549) xenograft mouse models, the loss of RAVER1 even reduced tumor growth, indicating that RAVER1 serves a rather pro-oncogenic role in lung and presumably other cancers. PTBPs and RAVER1 associate and co-localize in perinucleolar compartments (PNCs, (17)), which show elevated prevalence in progressed malignancies (22,23). Thus, it was tempting to speculate that RAVER1 modulates tumor cell fate in a PNC-dependent manner. However, in lung cancer cells, PNC number and integrity was not strictly associated with RAVER1 abundance but rather varied due to clonal variability. This indicates that RAVER1 is not essential for PNC assembly and controls tumor cell fate largely independent of PNC prevalence. The characterization of deregulated gene expression upon RAVER1 downregulation confirmed the observed defects in proliferation and apoptosis but we could not reveal an enrichment of RAVER1-guided AS in the respective pathways, e.g. E2F-driven gene expression. Most unexpectedly, however, RAVER1 depletion elevated TGFB signaling, epithelial-mesenchymal-transition (EMT) and strongly enhanced lethal EMT. The latter was proposed to promote tumor cell death upon exposure to elevated TGFB levels in a SNAIL/SOX4/KLF5-as well as BIM/BMF-dependent manner (36,37). Consistently, RAVER1 but not PTBP1 depletion elevated BIM, BMF, SNAI2 and SOX4 but decreased KLF5 even in the absence of TGFB exposure. This suggests that RAVER1 limits TGFB/BMP signaling probably by restricting TGFBR1/2 abundance.

The control of cancerous EMT essentially relies on the upregulation of central TGFB signaling factors, including TGFBR1/2, and enhanced expression of the EMT-driving transcription factor families SNAIs, TWISTs and ZEBs (38). Pivotal inhibitors of this process are microRNAs, in particular the miR-200 family, which impairs the synthesis of TGFB signaling components, including TGFBRs, as well as EMT-driving transcription factors (40,41). Our studies demonstrate that RAVER1 is a broad modulator of miR/RISC activity, but barely affects miR biogenesis. This regulatory role is associated with a substantial number of miR/RISC pathway genes subjected to control of AS by RAVER1. The multitude of splicing events modulated by RAVER1 within the miR/RISC pathway, affecting 20 of 29 major pathway genes, in conjunction with disturbed expression of some components, e.g. TNRC6 proteins, prohibits the identification of prime events via which RAVER1 modulates miR/RISC activity. However, evaluation of RAVER1-modualted splicing events in TNRC6 proteins provide strong evidence for a RAVER1-dependent modulation of protein interactions guiding miR/RISC activity like TNRC6-AGO2 association via GW-enriched regions in TNRC6 proteins (49). The fine-tuning of these interactions by shifting the isoform composition and abundance of miR/RISC pathway factors likely influences the activity and cooperativity of miR/RISC complexes, in a potentially mRNA-dependent but generally broad manner (50). In addition, RAVER1 likely influences the nuclear as well as cytoplasmic functions of the CCR4/NOT complex (51). RAVER1 associates with CCR4/NOT components, especially the scaffolding factor CNOT1, and extensively modulates splicing of most CCR4/NOT factors. How this influences CCR4/NOT functions and if this also provides a rationale for the potentially mRNA-specific modulation of miR-in/dependent deadenylation (52), requires further investigation.

In addition to providing the first mechanistic insights into how RAVER1 controls tumor cell fate, our studies also present the first transcriptome-wide investigation of the RAVER1-dependent control of splicing. So far, RAVER1 primarily was considered to serve as a co-factor of PTBP-dependent splicing (5,16,19). Our findings, confirm this synergy of PTBP1 and RAVER1 in controlling more than 1000 AS events in A549 cells. However, we also observe that RAVER1 regulates even more AS events (> 2500) largely independent of PTBP1. This suggests that RAVER1 also serves as a co-factor for additional splicing regulators, as supported by the enrichment of mRNA splicing factors among proteins co-purified with RAVER1 from nuclear fractions. Whether co-regulation with PTBP1 or other splicing factors involves direct association of RAVER1 with target transcripts remains to be determined. Attempts to evaluate RAVER1-RNA association by STAMP (Surveying Targets by APOBEC-Mediated Profiling, (53)) so far failed to reveal preferred binding substrates of RAVER1 (data not shown). This favors the view that RAVER1 mainly acts as a co-factor orchestrating the assembly of regulatory protein complexes guiding AS control, as already proposed for its synergistic regulation with PTBPs (18). The assembly of such multi-protein complexes on pre-mRNAs likely settles on RAVER1’s extended IDR, which is essential for its association with PTBPs. By acting in concert with such factors, RAVER1 likely modulates a multitude of AS events affecting tumor cell fate and the innate immune response (IIR). This was proposed due to RAVER1’s association with MDA5 and roles in modulating SARS-CoV-2 infections in human and primate cell models (54,55). However, protein interactions and key splicing substrates implicated in RAVER1’s role in the IIR remain to be determined.

## DATA AVAILABILITY

RNA-sequencing data are deposited on GEO: GSE232714.

## SUPPLEMENTARY DATA

Supplementary Data are available at to *be indicated upon publication*.

## Acknowledgements

The authors thank the Core Facility Imaging (CFI) and core unit for NGS analyses of the Martin Luther University (MLU) for broad support with image and NGS data analysis.

## Author contributions

A.W. and S.H. designed the experiments. A.W., N.B., S.M., KB.O., M.L., C.M., T.F., and C.I. performed experimental studies. M.G., L.S., and S.H. supported data analysis and interpretation. M.K., T.G., and A.S. supported experimental design and data interpretation. S.H. conceived the experimental design and wrote the manuscript.

Schemes were created with <u>BioRender.com</u>.

