## Supplementary Figures and Tables for "RAVER1 interconnects lethal EMT and miR/RISC activity by the control of alternative splicing": Supplementary Figures_Wedler et al_2023.pdf

**Supplementary Figure S1.** RAVER1 promotes tumor cell proliferation. **(A)** Scatter plot of RAVER1 and PTBP1 expression across all TCGA tumor cohorts (n=33). **(B)** Cell cycle progression analyses of indicated knockdowns in ES-2 and Hep-G2 cells, as described in Figure 1F.

**Supplementary Figure S2.** RAVER1 depletion promotes EMT. Immunostaining of CHD2, CTNNB1 and F-actin using Phalloidin-TRITC upon RAVER1- or PTBP1 depletion in A549 cells (left). Scheme of distinct adhesion junctions (AJ) morphology is shown in the right panel.

**Supplementary Figure S3.** RAVER1 modulates alternative splicing in synergy with and independent of PTBP1. **(A)** Heatmap depicting changes in miR/RISC pathway gene expression upon RAVER1 (siR) depletion in A549 cells, as described in Figure 4B. **(B-D)** Analysis of alternative splicing (AS) events and hexamer frequency upon PTBP1 depletion in A549 cells, as described in Figure 6B, C, and E. **(E)** Gene ontology (GO) analyses of shared splicing events, RAVER1_only and PTBP1_only according to Figure 6D. **(F)** Western blot analysis of SBP-Immunoprecipitation from nuclear extracts of RAVER1 (WT) for LC-MS/MS analyses described in Figure 6F.

**Supplementary Figure S4.** RAVER1 is a master regulator of alternative splicing within the miR/RISC pathway. **(A)** Analysis of selected alternative splicing (AS) events of the miR/RISC pathway by PCR as described in Figure 7C. Left panel shows semi-quantitative PCR for indicated genes, middle panel depict quantification of PCRs, and right panels provide schemes of extracted protein regions and domains along with alternatively spliced exons.

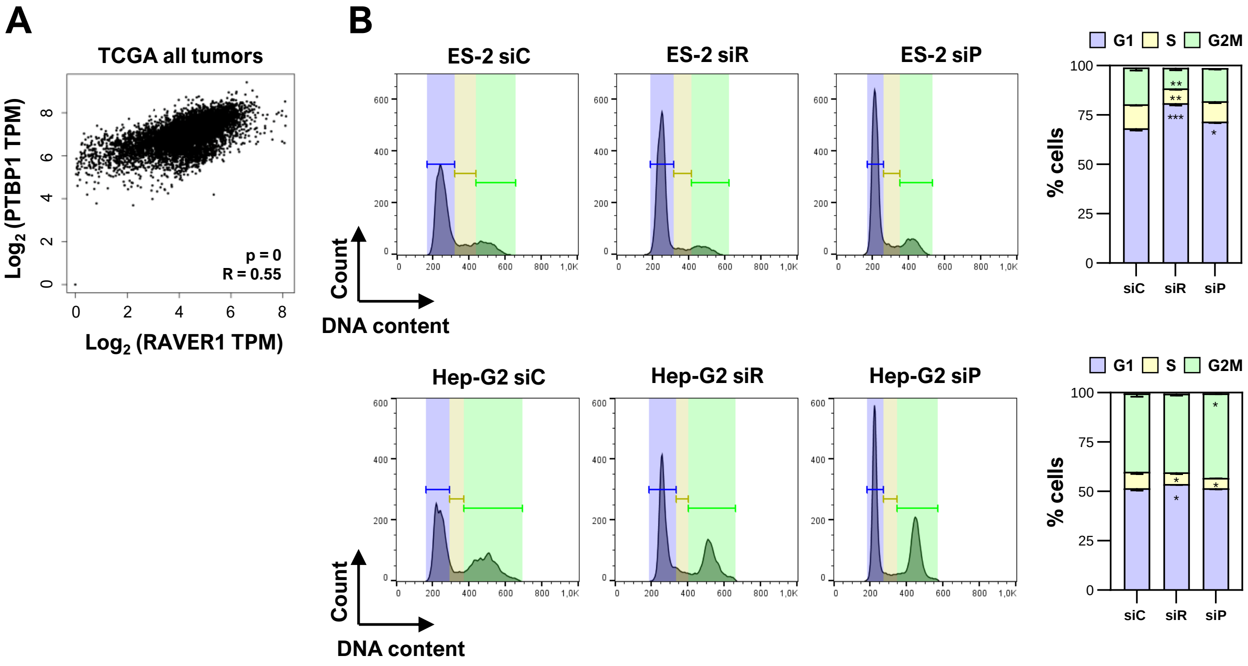


Supplementary Figure 1; Wedler et al., 2023

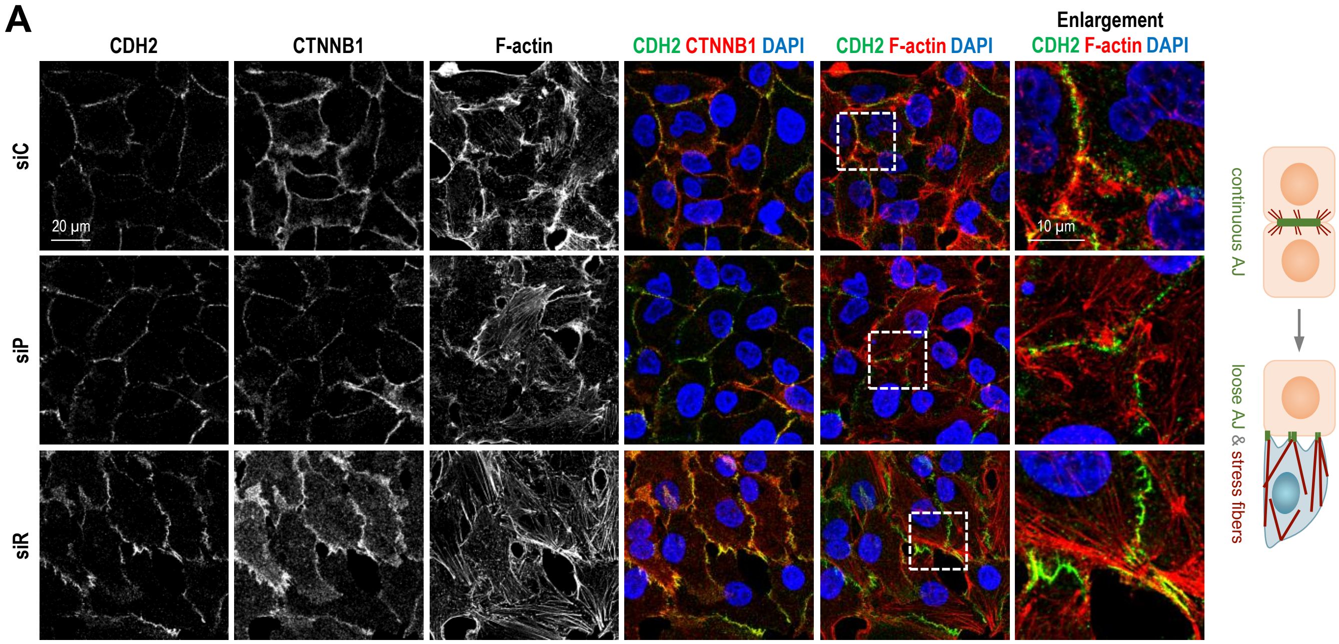


Supplementary Figure 2; Wedler et al., 2023

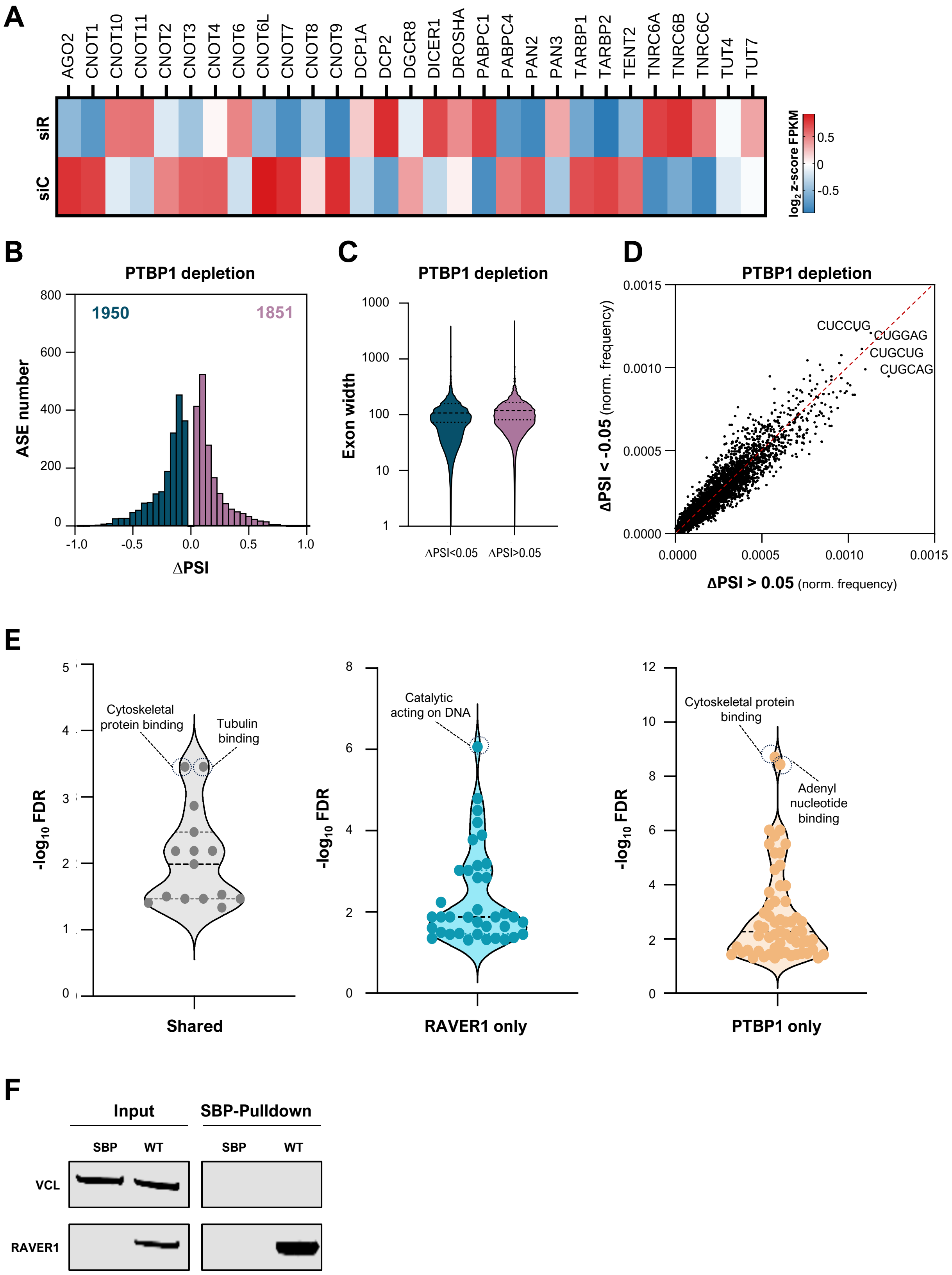


Supplementary Figure 3; Wedler et al., 2023

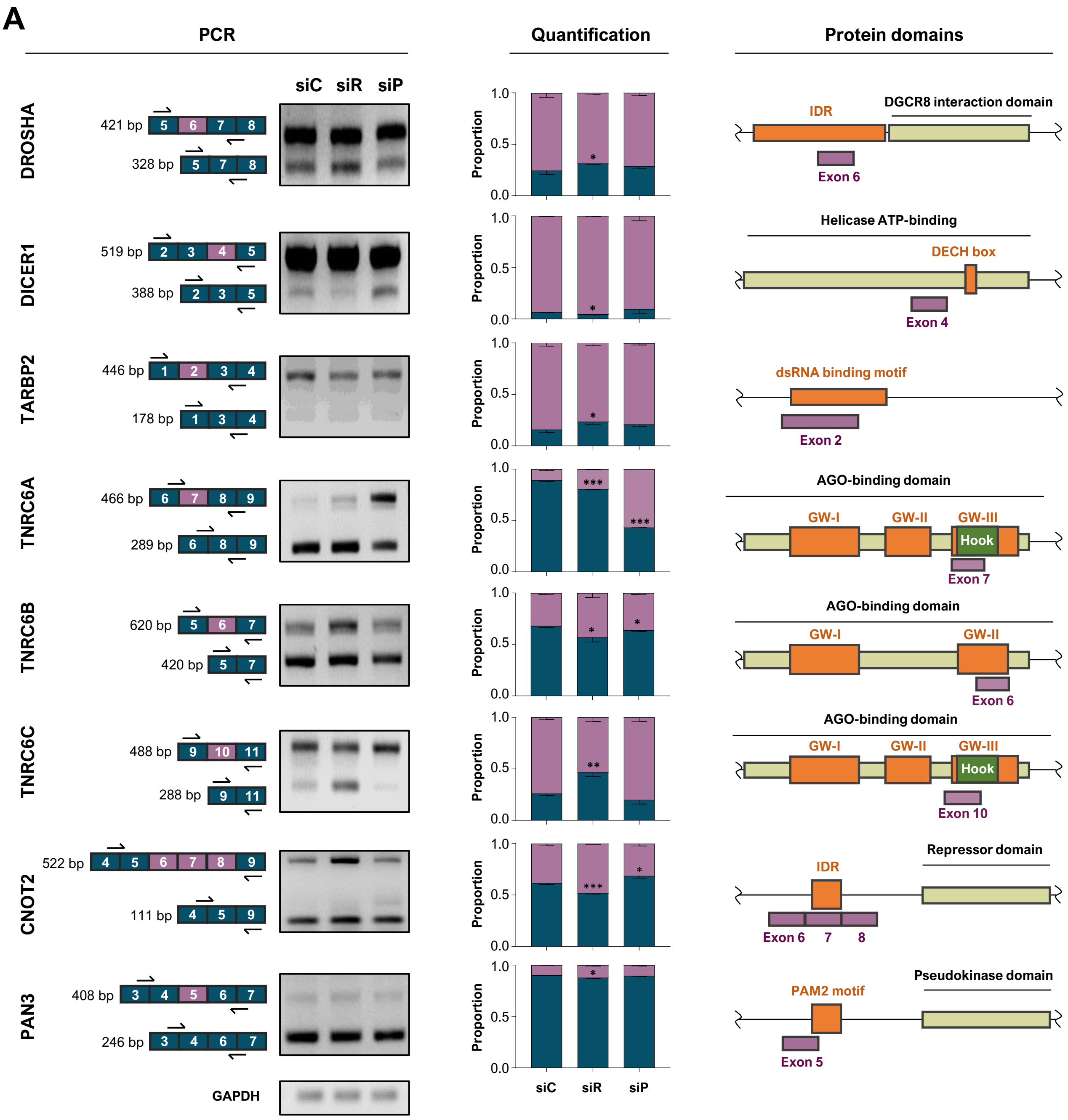


Supplementary Figure 4; Wedler et al., 2023
